# Aberrant DJ-1 expression underlies L-type calcium channel hypoactivity in tuberous sclerosis complex and Alzheimer’s disease

**DOI:** 10.1101/2020.08.26.267260

**Authors:** Farr Niere, Ayse Uneri, Colin J. McArdle, Luisa P. Cacheaux, Hailey X. Egido-Betancourt, Sanjeev Namjoshi, Cameron Reynoldson, Juan Penaranda, Xin Wang, William C. Taylor, Suzanne Craft, Christopher Dirk Keene, Tao Ma, Kimberly F. Raab-Graham

## Abstract

L-type voltage-dependent Ca^2+^ channels (L-VDCC) integrate synaptic signals to facilitate a plethora of cellular mechanisms. L-VDCC dysfunction is implicated in several neurological and psychiatric diseases. Despite their importance, signals upstream of L-VDCC activity that regulate their channel density, however, are poorly defined. In disease models with overactive mammalian target of rapamycin complex 1 (mTORC1) signaling (or mTORopathies), including tuberous sclerosis (TS) and Alzheimer’s disease (AD), we report a novel mechanism downstream of mTORC1 signaling that results in a deficit in dendritic L-VDCC activity. Deficits in L-VDCC activity are associated with increased expression of the mTORC1-regulated RNA-binding protein DJ-1. DJ-1 binds the mRNA coding the auxiliary Ca^2+^ channel subunit α2δ2 responsible for shuttling L-VDCC to the membrane and represses its expression. Moreover, this novel DJ-1/α2δ2/L-VDCC pathway is disrupted in human AD and preclinical models of AD and TS. Our discovery that DJ-1 directs L-VDCC activity and L-VDCC-associated protein α2δ2 at the synapse suggests that DJ-1/α2δ2/L-VDCC is a common, fundamental pathway disrupted in TS and AD that can be targeted in clinical mTORopathies.

**Significance Statement:** Many neurological disorders share symptoms, despite disparity among diseases. Treatments are prescribed based on diagnosis rather than individual symptoms. While only treating symptoms may obscure the disease, mechanism-based drug development allows the two approaches to converge. Hub proteins, those that coordinate the expression of proteins that mediate specific cellular functions, may be dysregulated across a broad range of disorders. Herein, we show that the RNA-binding protein DJ-1 controls the activity of L-type voltage-dependent calcium channels (L-VDCC), via the expression of its auxiliary subunit alpha2delta2 (α2δ2). Importantly, we demonstrate that this novel DJ-1/α2δ2/L-VDCC pathway is commonly disrupted among neurological disorders, namely Alzheimer’s disease (AD) and Tuberous Sclerosis (TS). Collectively, these data rationalize mechanism-based drug therapy to treat disease.

## Introduction

The mammalian target of rapamycin (mTOR) signaling pathway regulates the expression of proteins. Neurological disorders that manifest overactive mTOR (mTORopathy) are characterized by neuronal hyperexcitability and seizures, a shared symptom of autism spectrum disorder (ASD) and late-stage Alzheimer’s disease (AD) (1-4). The best example of a prototypical mTORopathy is tuberous sclerosis complex (TS). Individuals with TS display hyperactive mTOR signaling which is believed to be the root cause of epilepsy and ASD often associated with this disorder (5, 6).

Loss of function of either TSC1 or TSC2 protein increases mTOR signaling (7). mTOR forms two complexes—mTORC1 and C2—and it is well-established that the mTORC1 pathway promotes protein synthesis (8). It was therefore unexpected that a mouse model of TS exhibited reduced protein synthesis (9). Subsequent to this study, AD has also been found to exhibit reduced protein synthesis (10). Since then, compelling questions have arisen but remain unanswered: what mRNAs are repressed in mTORopathies, how are these mRNAs translationally regulated, and what are the functional consequences of repressing these mRNAs? Here, we demonstrate that DJ-1 (also known as Parkinsonism associated deglycase or Parkinson disease (autosomal recessive, early onset) 7 (Park7)), an RNA-binding protein, is a major translational hub disrupted in TS and AD. DJ-1 represses the translation of the calcium (Ca^2+^) channel subunit *Cacna2d2* mRNA, which encodes α2δ2 protein that facilitates the stability and trafficking of Ca^2+^ channels. Thus, elevated DJ-1 levels reduce the dendritic expression of α2δ2 and L-type voltage-dependent Ca^2+^ channel (VDCC) activity. These studies bring to light a novel mechanism of controlling local Ca^2+^ signaling that is commonly dysregulated in developmental-and aging-associated mTORopathies.

## Results

### DJ-1 target identification

We previously found that DJ-1 is associated with translation regulation and its expression is sensitive to mTORC1 activity in the postsynaptic density (PSD) isolated from the cortex (11). To test the role of DJ-1 as a translational hub that may be disrupted in mTORopathies, we first computationally defined the DJ-1-binding sequence (12). We found that the RNA sequences GNGCNG and CNGCNG are highly represented in known DJ-1-associated mRNAs (Fig. 1A) (12). Because DJ-1 is in the PSD, we sought the number of occurrences of these two DJ-1-binding motifs in mRNAs that encode PSD-associated proteins (11, 13). These mRNAs were ranked by frequency of DJ-1-binding sequence per kilobase (freq/kB). Of the 916 PSD-associated mRNAs, the mean freq/kB of both motifs in different mRNA regions are: 1.72 in 3’ untranslated region (UTR), 15.66 in cDNA or coding region (CR), and 0.43 in 5’UTR. The distribution of DJ-1-binding motifs throughout the mRNA suggests that DJ-1 may control the elongation step of protein synthesis, consistent with the role that DJ-1 represses its target mRNAs (12, 14, 15).

**Figure 1.**
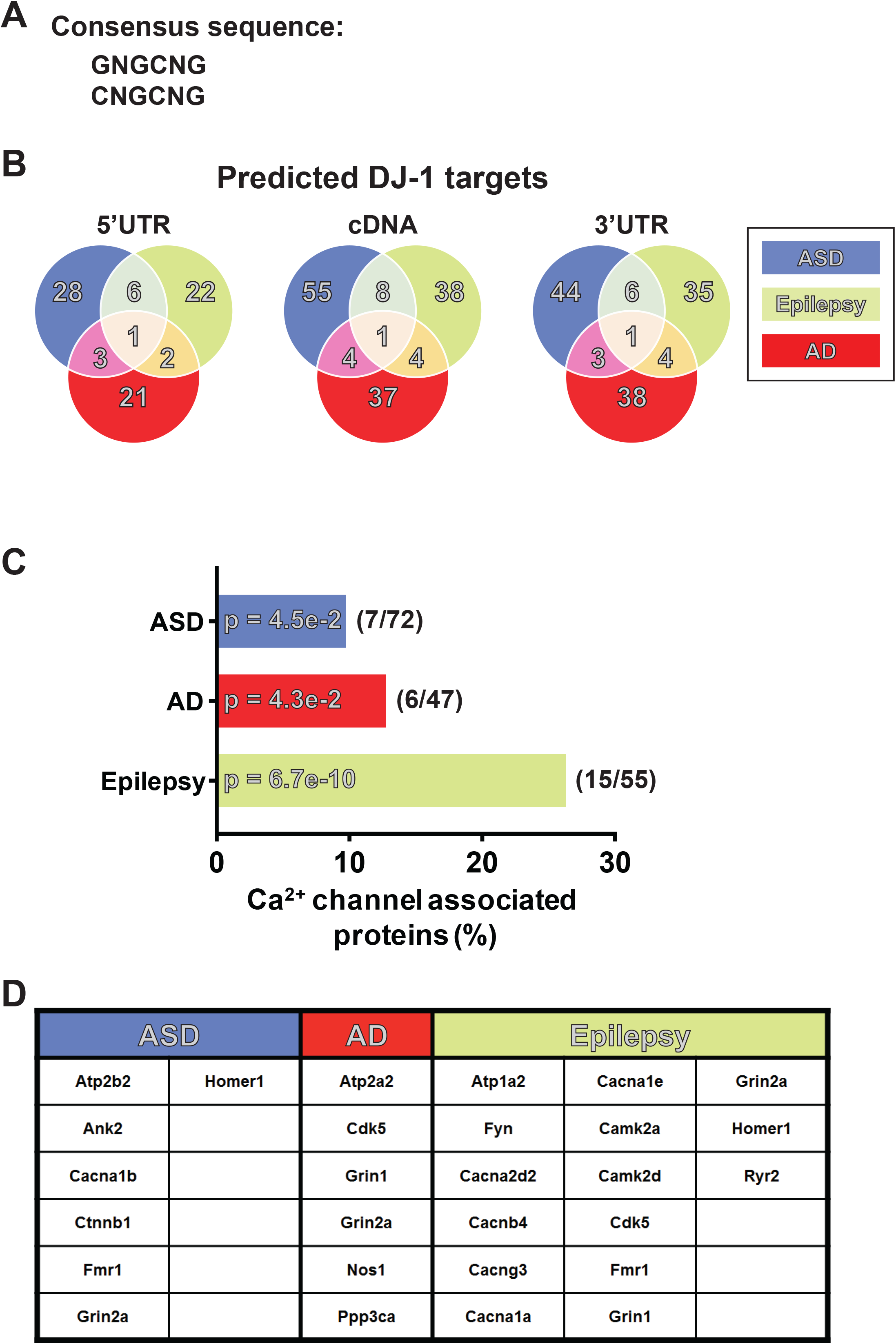
Autism spectrum disorder (ASD)-, epilepsy-, and Alzheimer’s disease (AD)- risk mRNAs contain putative DJ-1-binding motifs. **(A)** Putative DJ-1-binding sequence. **(B)** Venn diagram depicting DJ-1 target mRNAs whose proteins associate with the postsynaptic density (PSD). The mRNAs are grouped based on (1) the location of DJ-1- binding motifs in the sequence, namely 5’ untranslated region (UTR), coding region (cDNA) or 3’UTR, and (2) their linkage to ASD, epilepsy, and/or AD. **(C)** Percentages of risk mRNAs for ASD, AD, and epilepsy that encode Ca^2+^ channel-associated proteins and contain DJ-1-binding motifs. 10, 13 and 27 percent of DJ-1-target mRNAs linked to ASD, AD and epilepsy, respectively, participate in “Ca^2+^ ion transport” (Table S2). **(D)** List of DJ-1-target mRNAs whose proteins are implicated in “Ca^2+^ ion transport”.

DJ-1, prior to our report, has been primarily linked to Parkinson’s disease, yet never to epilepsy and ASD—symptoms that are common in TS—or AD (11). Recently, however, it has been identified as a potential biomarker for dementia (16). We reasoned, therefore, that DJ-1’s RNA-binding properties and target mRNAs may underlie a common pathology in these disorders. A Gene Ontology (GO) enrichment analysis of putative DJ-1 target mRNAs that encode PSD-associated proteins linked to epilepsy, AD, and ASD reveals that many of these targets are involved in ion transport (Table S1 and S2). Consistent with ion transport in the PSD, *Grin2a*, which codes for an ionotropic glutamate receptor, is the only DJ-1-target ion channel implicated in epilepsy, AD, and ASD. (Fig. 1B; Table S2, S3). Since NMDA receptor expression and function have been extensively characterized in these disorders, we decided to focus on DJ-1 targets related to epilepsy since seizures have been suggested to underlie both AD and ASD (17-22). We noted several coding for Ca^2+^ channels in epilepsy but not in AD or ASD. GO analysis of these mRNAs by disease indeed reveals an enrichment of “calcium ion transport”-associated proteins in epilepsy (Fig. 1C and D; Table S2). We focused on Ca^2+^-associated channels, because Ca^2+^ is arguably the most important second messenger in the brain as it is essential in neurodevelopment, synaptic transmission, and plasticity (23, 24). Moreover, disrupted Ca^2+^ homeostasis is suggested to be a crucial turning point leading to the progression of many neurodegenerative diseases (25). We focused on *Cacna2d2*, which encodes α2δ2, since α2δ2 promotes the forward trafficking of several pore forming Ca^2+^ channel subunits to increase current density and influence channel activation and inactivation kinetics (26-28). Additionally, loss of function mutations in CACNA2D2 gene can lead to seizure activity, a common symptom in epilepsy, ASD and AD (29). α2δ subunits are typically localized to presynaptic terminals; however, little is known about the subcellular localization of α2δ2 (30). *In situ* hybridization of *Cacna2d2* mRNA surprisingly indicates that it resides in the dendrites—in addition to the soma—intimating that α2δ2 can be synthesized upon changes in synaptic activity (Fig. 2A).

**Figure 2.**
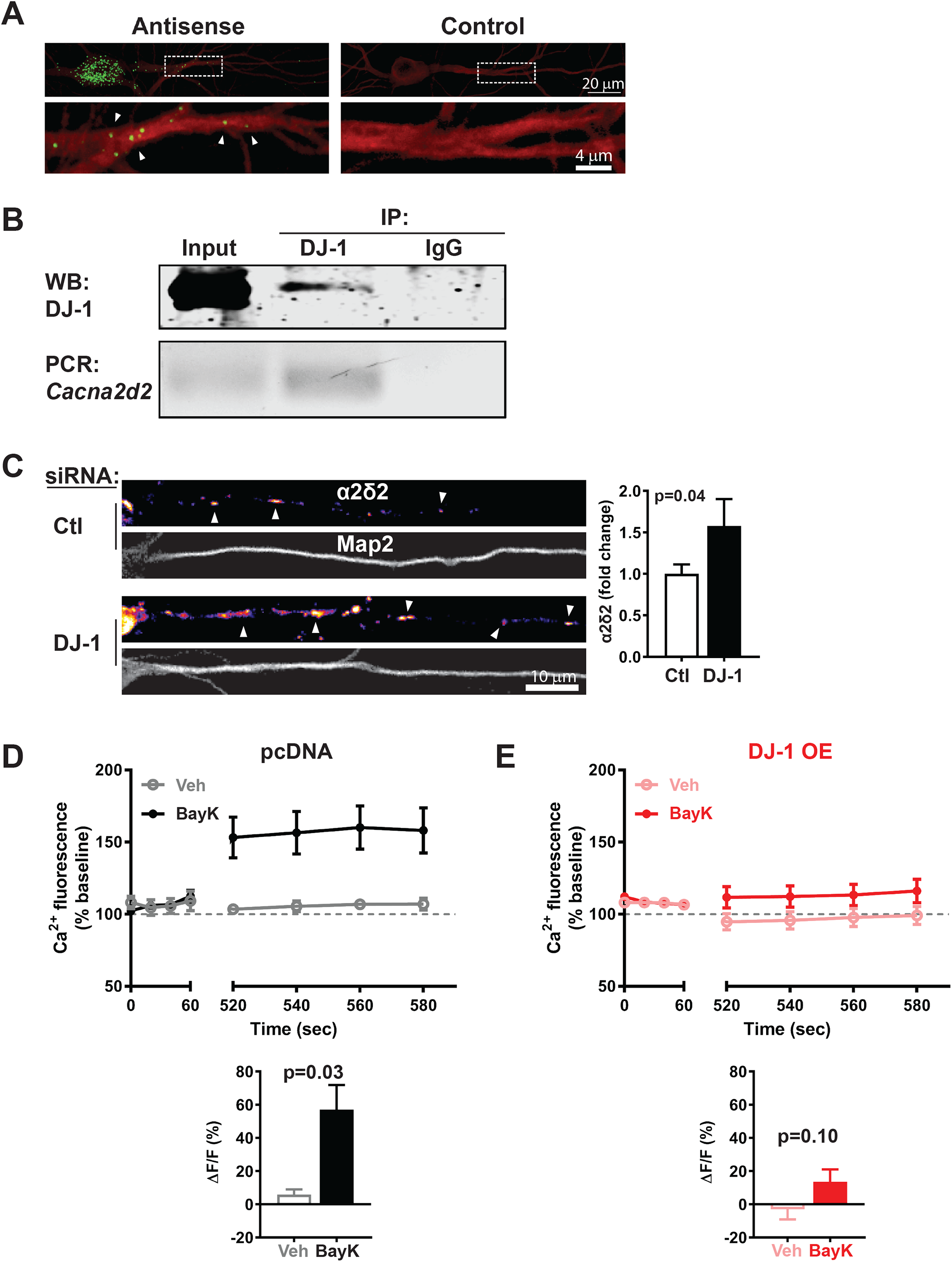
DJ-1 regulates the dendritic expression of α2δ2 and activity of L-type VDCC. **(A-C)** Characterization of *Cacna2d2* mRNA, which encodes α2δ2, as a DJ-1 target. **A**. (*top*) Representative fluorescence *in situ* hybridization of *Cacna2d2* mRNA depicting its presence in soma and dendrites. (*Bottom*) Magnified images of dendrites (outlined by broken lines) from top panel containing *Cacna2d2* mRNA (green dots/white arrowheads). **(B)** Co-immunoprecipitation of DJ-1 and *Cacna2d2* mRNA. (*Top*) Representative immunoblot of DJ-1 coimmunoprecipitating with anti-DJ-1 antibody. (*Bottom*) RT-qPCR-amplified product of *Cacna2d2* mRNA isolated from DJ-1 immunoprecipitation. **(C)** Knockdown of DJ-1 increases α2δ2 protein. (*Left*) Representative images of α2δ2 protein (fire/white arrows) in neurons (*top*) expressing scrambled (control/Ctl) and (*bottom*) DJ-1/knockdown siRNAs. Map2 marks dendrites. (*Right*) Quantification of dendritic α2δ2 normalized to Ctl. Ctl: 1.00±0.11, n=23; DJ-1: 1.58±0.32 n=6. **(D and E)** Increased expression of DJ-1 inhibits L-type VDCC activity. Assessment of dendritic L-type Ca^2+^ channel activity in **(D)** pcDNA-expressing and **(E)** DJ-1-overexpressing dissociated hippocampal neurons using L-type agonist, BayK-644 (BayK, 5 μM). (*Top*) Average trace of Ca^2+^ fluorescence signal before (baseline; 0-60 seconds (sec)) and after (520-580 sec) addition of BayK or vehicle at ∼90 sec. (*Bottom*) Quantification of change in fluorescence (ΔF) normalized to baseline (F). **(D)** BayK, but not vehicle (Veh), increases Ca^2+^ fluorescence in pcDNA-expressing dendrites. Veh: 5.80±3.16, n=4; BayK: 57.07±14.81, n=6. **(E)** Overexpressing DJ-1 blocks BayK- induced increase in Ca^2+^ fluorescence. Veh: −3.10±6.04, n=17; BayK: 13.44±7.57, n=11. Values are shown as mean±s.e.m.; statistics: student’s t-test.

*Cacna2d2* mRNA has several predicted DJ-1-binding sites (5’UTR: 15-26; CR: 58-61; 3’UTR: 7-8) (Table S3). To test if *Cacna2d2* mRNA is a DJ-1 target, we performed RNA immunoprecipitation (RNA-IP) by isolating both DJ-1 protein and *Cacna2d2* mRNA from anti-DJ1 RNA-IP fraction. Western blot analysis demonstrates that DJ-1 protein binds to DJ-1 antibody (Fig. 2B). Furthermore, reverse-transcription followed by semi-quantitative polymerase chain reaction (RT-qPCR) detects *Cacna2d2* mRNA specifically in DJ-1 and not the negative IgG control (Fig. 2B). These data confirm that *Cacna2d2* mRNA is a target of DJ-1. To determine if DJ-1 promotes or represses *Cacna2d2* mRNA translation, we knocked down DJ-1 in cultured hippocampal neurons by siRNA and measured α2δ2 protein levels by immunocytochemistry. Knockdown of DJ-1 increases α2δ2 in dendrites by ∼60%, confirming a role for DJ-1 as a translational repressor (Fig. 2C) (12).

### DJ-1 attenuates L-VDCC activity

α2δ subunits associate with voltage-dependent Ca^2+^ channels (VDCC) (26-28). Since perturbing DJ-1 levels alter α2δ2 expression in the dendrites, we predicted that DJ-1 can also affect a dendritic VDCC through α2δ2 levels. Notably, L-type VDCC, which is open at rest, is prominently expressed on dendrites (31). We, therefore, assessed L-type channel function by performing live Ca^2+^ imaging in dissociated rat hippocampal neurons that overexpress DJ-1. We activated L-type VDCC with its channel opener, BayK-8644 (BayK, 5 μM) and measured the change in dendritic Ca^2+^ fluorescence (ΔF/F). Application of BayK on pcDNA-expressing neurons, which serve as control, increases dendritic fluorescence by ∼30% (Fig. 2D). Interestingly, DJ-1-overexpressing cells fail to display BayK-induced elevation of Ca^2+^ signal in dendrites (Fig 2E). These data link DJ-1 to dendritic L-type VDCC function and perhaps the signaling downstream of L-type channels. Through bioinformatics analyses and experimental validation, these data collectively establish that DJ-1 can regulate Ca^2+^ dynamics through α2δ2 expression and L-type VDCC activity.

### Aberrant α2δ2 expression and L-VDCC activity in TS

Since dendritic DJ-1 is excessively expressed in a preclinical mouse model of TS, we hypothesized that TS and other mTORopathies have disrupted Ca^2+^ channel activity (11). We first measured α2δ2 levels, *in vivo* and *in vitro*, after knocking out the *Tsc1* gene by Cre expression (*Tsc1 KO* or TS) in dissociated hippocampal neurons and hippocampal slices. Immunostaining for α2δ2 indeed shows that TS neurons have ∼30% lower α2δ2 levels in the dendritic fields of hippocampal CA1 region and in dendrites of cultured hippocampal neurons compared to control (Fig. 3A and B). Low α2δ2 levels predict reduced Ca^2+^ signaling in TS dendrites (27). To determine if TS dendrites have reduced L-type currents, we activated L-type VDCC with BayK and measured dendritic ΔF/F. WT dendrites, as expected, show a ∼25% increase in ΔF/F with BayK (Fig. 3C) (26, 27, 30). Surprisingly, BayK-induced L-type activity is absent in TS, similar to DJ-1 overexpression (Fig. 2E). These data bring to light that TS harbors abnormal L-type VDCC activity, a newly-identified TS-associated dysfunction.

**Figure 3.**
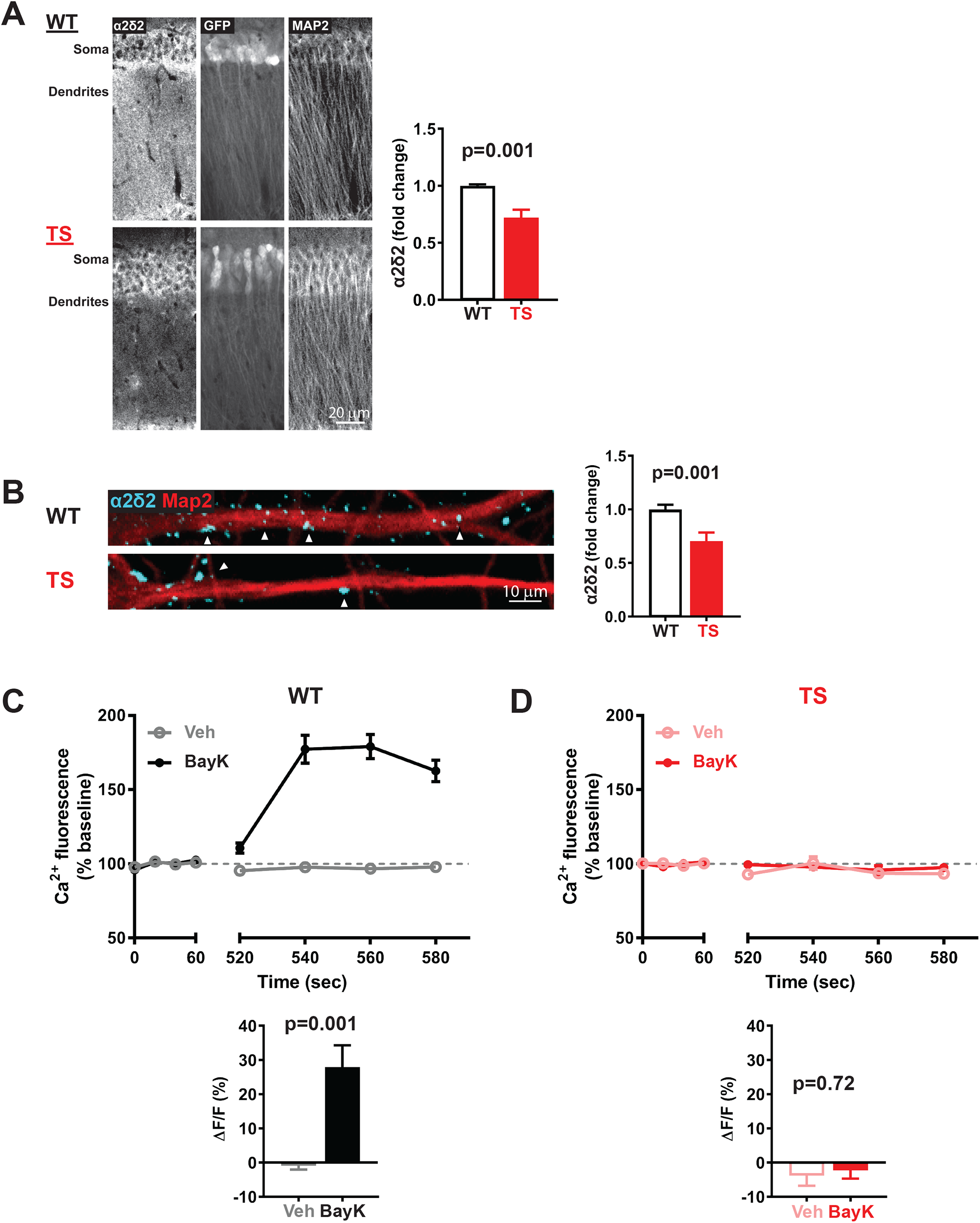
Preclinical models of TS exhibit diminished α2δ2 expression and L-type VDCC activity. **(A)** (*Top*) Compared to wildtype mice expressing GFP alone (WT), (*bottom*) Cre-GFP-mediated knockdown of *Tsc1* (TS) in the hippocampus reduces basal α2δ2 levels in *stratum radiatum* of CA1 area as shown by immunohistochemistry of acute slices. GFP indicates Cre expression and Map2 marks dendrites. Quantification normalized to WT (WT: 1.00±0.01, n=12; TS: 0.72±0.07 n=12). **(B)** (*Top*) Basal expression of α2δ2 in dendrites of WT dissociated hippocampal neurons is significantly higher than (*bottom)* that of Cre-GFP-expressing neurons (TS) as shown by immunocytochemistry. Map2 marks dendrites. Quantification normalized to WT (WT: 1.00±0.04, n=37; TS: 0.70±0.08 n=19). **(C and D)** L-type VDCC activity is absent in TS. **(C)** While activation of L-type channels by BayK increases Ca^2+^ fluorescence in WT dendrites (Veh: −0.96±1.12, n=20; BayK: 27.92±6.39, n=14), **(D)** BayK fails to increase Ca^2+^ signal in TS (Veh: −3.82±2.98, n=28; BayK: −2.29±2.44, n=18). Values are shown as mean±s.e.m.; statistics: student’s t-test

### Aberrant α2δ2 and CaV1.2 expression and L-VDCC activity in AD

Do aberrant DJ-1 and α2δ2 levels and L-type channel signaling constitute a molecular signature of a mTORopathy? To answer this question, we used a preclinical mouse model of Alzheimer’s disease—APP/PS1 transgenic mouse—since aberrant mTOR signaling is observed in AD (2-4). We prepared dissociated hippocampal neurons from APP/PS1 mice and their wildtype littermates. Similar to our preclinical TS mouse model, hippocampal neurons of APP/PS1 mice have ∼200% more dendritic DJ-1 and ∼50% less α2δ2 levels compared to wildtype controls (Fig. 4A and B). Moreover, BayK-induced L-type signaling is attenuated in APP/PS1 dendrites (Fig. 4C and D). In agreement with our prediction, the preclinical mouse model of AD harbors similar abnormal molecular signature (DJ-1/α2δ2/L-VDCC) as in the TS mouse model.

**Figure 4.**
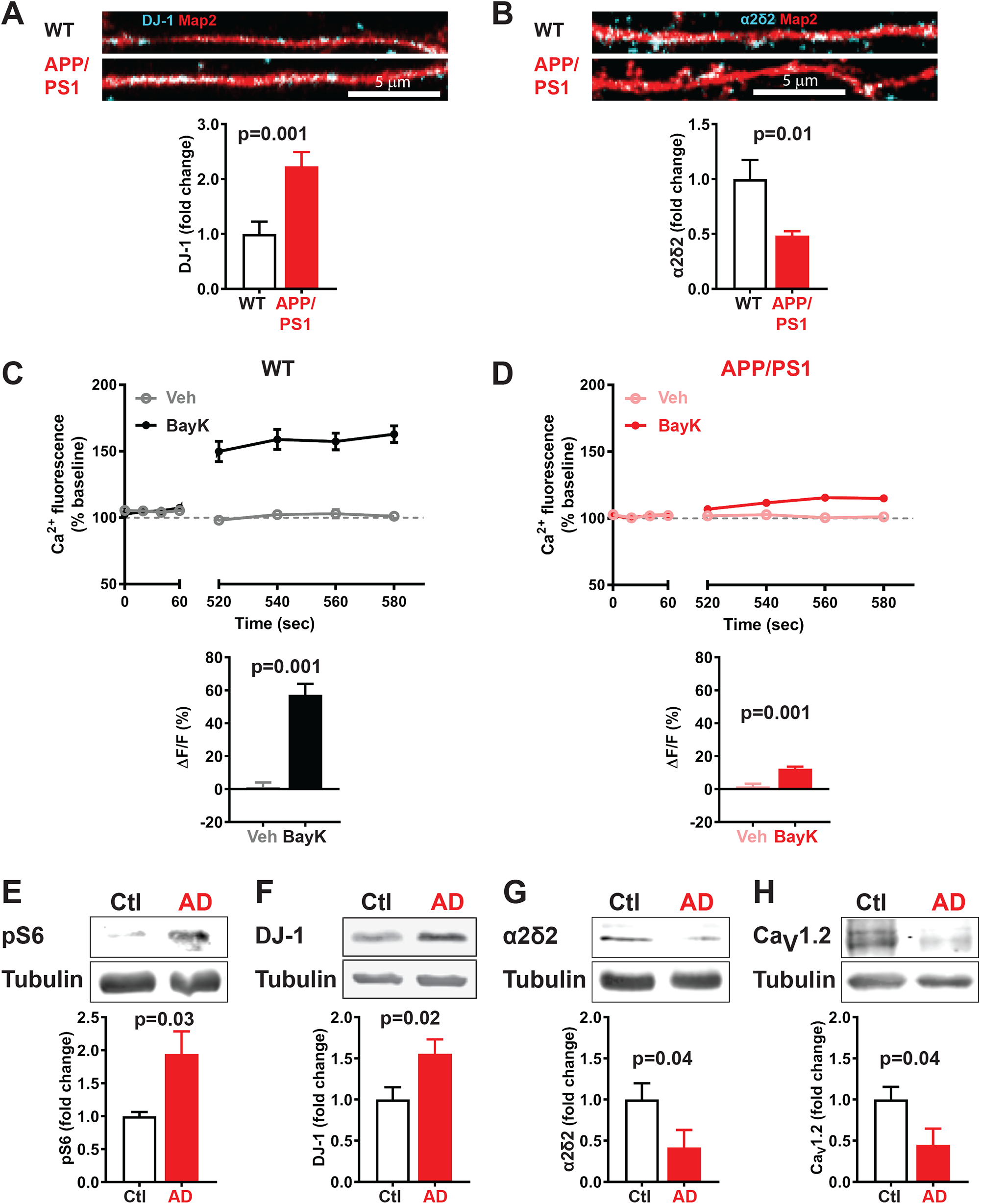
The APP/PS1 preclinical model of and human brains with AD exhibit disrupted DJ-1 and L-type-VDCC-associated protein expression and diminished L-type VDCC activity. **(A and B)** Immunocytochemical analyses in dissociated hippocampal neurons from littermates expressing normal (WT; *top image*) and mutant APP and PS1 genes (APP/PS1; *bottom image*) demonstrate that **(A)** DJ-1 expression (blue) in WT dendrites is markedly lower than APP/PS1 as quantified and normalized to WT (WT: 1.00±0.23, n=27; APP/PS1: 2.24±0.26, n=30; p<0.001); and **(B)** α2δ2 level (blue) in WT dendrites is higher than APP/PS1, as quantified and normalized to WT (WT: 1.00±0.17, n=30; APP/PS1: 0.48±0.04, n=30). **(C and D)** L-type Ca^2+^ channel activity is attenuated in APP/PS1 model of AD. **(C)** In dissociated WT hippocampal neurons, BayK but not vehicle increases dendritic Ca^2+^ fluorescence. Veh: 1.22±2.81, n=6; BayK: 57.29±6.65, n=5. **(D)** Neurons from APP/PS1 littermates display reduced Ca^2+^ fluorescence in dendrites with BayK. Veh: 1.60±1.66, n=8; BayK: 12.30±1.29, n=7. **(E-H)** Synaptoneurosomes of human brains diagnosed with AD present mTOR- and L-VDCC- associated protein changes similar to mouse models of TS and AD. (*Top*) Representative western blots and (*bottom*) quantification are shown. **(E-F)** Compared and normalized to age-matched controls (Ctl), AD samples have overactive mTOR as measured by elevated **(E)** phospho-S6 (Ctl: 1.00±0.07, n=3; AD: 1.94±0.34, n=3) and **(F)** DJ-1 (Ctl: 1.00±0.15, n=5; AD: 1.56±0.18, n=5) immunoblots. **(G-H)** L-VDCC activity may be attenuated in AD since the following levels of L-type-associated proteins are diminished in AD: (G) α2δ2 (Ctl: 1.00±0.20, n=5; AD: 0.42±0.21, n=5) and (H) L-VDCC pore-forming subunit CaV1.2 (1.00±0.15, n=4; AD: 0.45±0.20, n=3). Values are shown as mean±s.e.m.; statistics: student’s t-test.

As preclinical models of TS and AD show the same abnormal molecular signature—elevated DJ-1 and diminished α2δ2 levels and L-type VDCC hypoactivity— we sought whether these signs are recapitulated in human AD. Using synaptoneurosomes isolated from human prefrontal cortices (PFC), we first measured mTOR activity in control and AD-diagnosed tissues. Subject gender, age, beta-amyloid load, Braak score, and dementia status are reported (Table S4). Western blot analyses of phospho-S6, a downstream marker of mTOR activity, is significantly elevated by ∼200% in AD samples, consistent with previous reports that mTOR is hyperactive in AD (Fig. 4E) (32-34). We next assessed DJ-1 expression as it is regulated by mTOR activity (11). DJ-1 expression, similar to TS and AD mouse models, is ∼50% more in AD compared to age-matched control samples (Fig. 4F). Because DJ-1 interacts with *Cacna2d2* mRNA (Fig. 2B) and abates dendritic α2δ2 levels (Fig. 2C), we measured α2δ2 in human PFC synaptoneurosomes. Recapitulating the observations in AD and TS mouse models, the human AD samples express ∼50% less α2δ2 levels than control (Fig. 4G). L-type VDCC is hypoactive in dendrites of preclinical models of TS and AD.

As a surrogate for L-type channel activity in postmortem human brain tissue, we hypothesized that the protein expression of CaV1.2—the pore-forming subunit of L-type Ca^2+^ channels—in AD samples is low compared to control. CaV1.2 expression is indeed ∼50% lower in human brains with AD than controls (Fig. 4H). Altogether, these data suggest that TS and AD share dysregulated L-type VDCC-associated protein expression that effect calcium signaling in dendrites.

## Discussion

As clinical trials, to date, have failed or produced mixed results in alleviating cognitive issues in both AD and TS, it is critical to determine the molecular mechanisms that underlie synaptic dysfunction in these disease (35). Here, using an unbiased, bioinformatics approach, to identify new ASD-and AD-related proteins. Notably, an enrichment of epilepsy-related Ca^2+^ channels that are DJ-1 target mRNAS, justified the exploration of Ca^2+^ dynamics in other mTORopathies with excessive expression of dendritic DJ-1. This finding also allowed us to uncover a novel L-VDCC pathology shared between TS and AD. Moreover, DJ-1 may impact several Ca^2+^ associated mRNAs, as predicted by our bioinformatics (Fig. 1) and Ca^2+^ dysregulation may be a common feature of mTORopathies. These discoveries may lead to the development of new mechanism-based treatments based on synaptic gene expression regulated by a single RBP *(see accompanying manuscript)*.

L-type VDCC has been suggested to be a homeostat for multiple pathways and dysregulated in ASD (36, 37). Our data expand the role of L-type VDCC as a central homeostat for mTORopathies beyond ASD. The animal models of TS and AD that we use here express elevated DJ-1, low α2δ2, and repressed L-type VDCC activity. L-type VDCCs are critical for normal synaptic function, and altering their activity is linked to neurological disorders, such as seizures, a core symptom of TS and AD (36, 38) *(see accompanying manuscript)*. Importantly, L-VDCC blockers are being considered as potential drug therapies (36-38). Our data, however, suggest that blocking L-VDCC may not be suitable for ASD, TS, AD, and perhaps for other mTORopathies, since L-VDCC function in these neurological disorders is already absent or low at best. Moreover, these data collectively may explain why L-VDCC blockade can facilitate subtle, nonconvulsive epilepsy, paralleling what is observed in TS and dementia patients (39-42). In summary, our work identifies the DJ-1/α2δ2/L-VDCC pathway in AD and TS, providing new druggable targets for the treatment of these disorders.

## Methods

### Determination of DJ-1-binding motifs and disease associated-gene overlap

Using a 45-nucleotide consensus sequence previously identified, we constructed a list of all possible 6-12 nucleotide substrings, a typical length of RNA-binding motifs (12, 43-45). Each substring was then ranked based on the sum of its probability values based on the probabilities derived from the consensus logo. We found that two motifs GNGCNG or CNGCNG occurred with highest probability. To find the frequency of occurrences for these motifs in a given DNA alphabet, we used the seqinr and Biostrings packages (46, 47). The graphical user interface for this process was developed using the shiny package in R along with the DT package (48, 49). PSD gene data were obtained from the BAYES-COLLINS-MOUSE-PSD-CONSENSUS available online at G2C: Genes to Cognition database (http://www.genes2cognition.org/) (13). All gene sequences were obtained using the biomart package (50-52). To create the Venn diagram, we first determined the frequency per kilobase (freq/kB) of putative DJ-1 RNA-binding motifs (GNGCNG or CNGCNG) within the 5’ UTR, cDNA, and 3’ UTR sequences of the 916 PSD genes with available sequence data. All matches were filtered to obtain only the maximum length sequences and then ranked by freq/kB in descending order. This list was then subset based on the overlap with mTOR-regulated proteins associated with epilepsy, Alzheimer’s disease, and autism spectrum disorder. Finally, the lists were thresholded using Grin2a as a cutoff and generated the Venn diagram. Disease-associated mTOR-regulated genes were obtained from databases as described by Niere *et al*, 2016 (11).

### Mice

#### Tsc1 conditional knockout

For the TSC mouse model, *Tsc1*^*tm1Djk*^/J mice (Jackson Laboratory, Bar Harbor, ME) were used (53). To generate Tsc1 KO, recombinant adeno-associated virus coding for Cre recombinase (AAV-Cre-GFP) was introduced in *Tsc1*^*tm1Djk*^/J mice to knockout the *tsc1* gene, and AAV-GFP (vector) served as control(11). Both AAV-GFP and AAV-CRE-GFP were driven by a synapsin promoter. As previously described for *in vivo* studies, AAV-Cre-GFP or AAV-GFP was injected into the hippocampus of 7 to 8 weeks-old male mice using the following coordinates (from bregma): ±2.2 mm A/P, ±1.5 mm M/L; ±2.5 mm A/P, and ±1.6 mm M/L (11). Two weeks after introducing one of the two viruses, hippocampal slices were generated and used for experiments. Male and female mice were used to generate dissociated hippocampal neurons.

#### APP/PS1

For the AD animal model, APP/PS1 mouse line which expresses human/murine amyloid precursor protein (APP) construct containing the Swedish double mutation (APPswe) and presenilin (PSEN1 or PS1) construct without the exon 9 (PSEN1/ΔE9) (2). Male and female mice were used, and WT littermates served as control for studies involving APP/PS1.

### Cell culture

Dissociated hippocampal neurons, 18–25 days *in vitro* (DIV), were prepared from embryonic rat pups (E17–18) similar to Sosanya *et al* (54). For *Tsc1* or APP/PS1 cultures, neurons were prepared from postnatal (0–3 days) *Tsc1* conditional knockout or APP/PS1 pups similar to Niere *et al* (55).Briefly, hippocampi were extracted from postnatal day 1–3 *Tsc1*^*tm1Djk*^/J or APP/PS1 mouse pups. The tissue was dissociated and plated in neurobasal A medium supplemented with B27, glutamine, and 1% fetal bovine serum. Cultures were plated at a density of ∼100,000 cells per 12 mm on glass coverslips that had been coated overnight with 50 μg ml^−1^ poly-D-lysine and 25 μg ml^−1^ laminin in borate buffer. Cultures were fed after 1 day after plating, and media was replaced approximately once a week with either fresh rat culture media (neurobasal A supplemented with B27, glutamine and 3 μM cytosine arabinoside (AraC)) or fresh mouse culture media (glial-conditioned media with 3 μM AraC) until cultures were used at DIV 14–21.

### Fluorescence in situ hybridization (FISH)

*Cacna2d2* mRNA detection was conducted using the ViewRNA ISH Cell Assay kit (Affymetrix) as described in Cajigas et al, 2012 (56). The *Cacna2d2* probe set was designed commercially by Affymetrix. Briefly, primary hippocampal neurons (days *in vitro* (DIV) 20-21) were fixed at room temperature for 30 minutes with a 4% paraformaldehyde solution (4% paraformaldehyde, 5.4% glucose, 0.01M sodium metaperiodate, in lysine-phosphate buffer). Proteinase K treatment was omitted and the rest of the hybridization was completed according to the manufacturer’s instructions. The cells were then washed with PBS and blocked with 4% goat serum in PBS for one hour followed by incubation in primary antibody (chicken anti-GFP) overnight at 4°C. After three washes with PBS the cells were incubated with the appropriate secondary antibody for one hour at room temperature and washed with PBS. The coverslips were then mounted with an antifading mounting medium and imaged as described above.

### RNA immunoprecipitation (RIP)

Mouse hippocampal tissue (pooled hippocampus from 6 mice) was homogenized in PLB buffer and incubated for 10 minutes on ice. The samples were then centrifuged for 14,000 xg for 15 minutes at 4°C. Protein concentration of the supernatant was determined with a BCA assay. The supernatant was then precleared with 5 μg goat IgG and 50 μl beads (Protein G) for 30 minutes at 4°C. Beads washed with NT2 buffer were incubated with either 30 μg DJ-1 antibody (Santa Cruz) or goat IgG for 2-3 hours at 4°C. The beads were washed 3 times with NT2 buffer. The precleared samples (4.5 mg) were incubated with the conjugated beads overnight at 4°C. The beads were washed 4 times with NT2 buffer and after the last wash were suspended in 200 μl NT2 buffer. 50 μl of the bead suspension was removed and incubated with 2X Laemmli sample buffer containing BME for 30 minutes at room temperature. The supernatant was then used for western blot analysis to confirm DJ-1 protein was immunoprecipitated. The rest of the bead suspension in NT2 buffer was used for RNA extraction. The supernatant was removed and RNA extraction was performed with TRIzol. A portion of the eluted RNA was used for quality assessment using an Agilent Bioanalyzer. Reverse transcription was performed with the iScript cDNA synthesis kit (Bio-Rad) according to the manufacturer’s instructions. PCR amplification was performed with commercially available gene specific primers for Akt and Cacna2d2 (GeneCopoeia) and PCR products were then run on a 2% agarose gel.

### Transfection

Primary hippocampal neurons were transiently transfected on DIV 14 with siRNA directed against DJ-1(SMARTpool: ON-TARGETplus DJ-1 siRNA, Cat No. L-050181-01-0005, Dharmacon) or scrambled siRNA (Dharmacon) plus a tdTomato plasmid; DJ-1 (pGEX-5X-1-DJ1-WT; Addgene) or pcDNA plus RFP plasmid(57) Hippocampal cultures were transfected with Lipofectamine 2000 (Invitrogen) according to the manufacturer’s instructions. Transfection was allowed to occur for 2.5 hours. Cells were processed for immunohistochemistry on DIV18.

### Immunohistochemistry (IHC)

For hippocampal slices, mice were transcardially perfused with phosphate buffered saline (0.1 M). Brains were removed and postfixed in 4% paraformaldehyde. 50 μm thick slices were prepared and checked for GFP expression. Slices expressing GFP were processed for IHC, similar to Niere *et al*, using the following primary antibodies: rabbit anti-α2δ2 (1:100, Novus), mouse anti-MAP2 (1:500, Abcam), and chicken anti-GFP (1:500, Aves) (55). With dissociated hippocampal neurons, cells were fixed in 4% paraformaldehyde for 20 minutes at room temperature and permeabilized in 0.2% Triton X-100 for 10 minutes (55). Fixed cells were incubated overnight in 4°C with the following primary antibodies: rabbit anti-α2δ2 (1:500, Alomone) or rabbit anti-DJ-1 (1:1000, Novus), and chicken anti-Map2 (1:1000, Aves) or mouse anti-Map2 (1:1000, Abcam). Appropriate secondary antibodies (1:500, Life Technologies) were used— AlexaFluor488 (AF488), AF647, AF405—after overnight primary antibody incubation.

### Calcium imaging

Dissociated hippocampal neurons at DIV 14–21 were used for live calcium imaging. Prior to imaging, cells were incubated in ACSF with Oregon Green 488 BAPTA-1 AM (OGB, 200 μM; 30 minutes; 37°C; ThermoFisher) as described by Workman *et al* (58). Cells were then transferred to fresh ACSF (37°C) for imaging (1 frame per 20 seconds). Baseline calcium signal was imaged (1 minute), after which (S)-(-)-Bay-K-8644 (5 μM, Tocris) or vehicle (DMSO) was added. Neurons were imaged for ∼600 s at room temperature. Quantification of the calcium signal was performed using Metamorph (Molecular Devices). Briefly, dendritic regions of interest (ROI) that were at least 5 μm from the soma were analyzed. The mean intensity values for each ROI at every 20 seconds were averaged as baseline (*F*_0_). The ROI intensity values obtained at each time point after the addition of BayK-8644 or vehicle were averaged (*F*). The equation, Δ*F*/*F*=((*F*−*F*_0_)/*F*_0_), was used to measure the change in signal and data were plotted as a percentage of the baseline (58).

### Western blots

#### Subjects and Tissues

Postmortem human prefrontal cortex (PFC) tissue samples were acquired from the University of Washington (UW) Neuropathology Core in accordance with the UW and Wake Forest University School of Medicine Institutional Review Boards. Tissues were harvested from patients that were clinically diagnosed with AD, where the neuropathology of AD was confirmed and from controls who exhibited low levels of AD neuropathology. The samples in this study underwent rapid autopsy shortly after death, where the harvested tissue was flash frozen. Clinical and neuropathological diagnoses were based on postmortem senile plaques, Braak neurofibrillary tangle staging and Consortium to Establish a Registry for Alzheimer’s Disease (CERAD) scores, collectively known as the “ABC score”. The patient characteristics are shown in Table S4. Mean age of death is 86.9 years, and the postmortem interval (PMI) ranged between 2 and 5 hours with an average of 4.03 hours.

#### Synaptoneurosome preparation

Frozen postmortem human prefrontal cortex samples from UW Neuropathology Core were homogenized with lysis buffer composed of Tris-Base with Halt™ Phosphatase and Protease Inhibitor Cocktail (Thermo Scientific). Homogenates were then sequentially filtered through 100μm cell strainer and 5μm syringe filter and centrifuged for 20 minutes in 4°C at 14,000 xg. The pellet was solubilized with radioimmunoprecipitation (RIPA) buffer and centrifuged for 10 minutes in 4°C at 14,000 xg. The supernatant was collected for protein quantification using Pierce™ BCA Protein Assay Kit (Thermo Scientific).

#### Western blot analysis

Samples were prepared into SDS-polyacrylamide gel electrophoresis (PAGE) loading buffer using 4x Laemmli Sample Buffer (Bio-Rad). 50 μg protein from each sample were resolved in 10% SDS-polyacrylamide gel and transferred onto 0.2 μm nitrocellulose membranes (Bio-Rad). The nitrocellulose membranes were blocked in 5% nonfat dry milk in Tris-buffered saline containing 0.1% Tween 20. To visualize the proteins, we used: mouse anti-DJ1 (1:2000; Novus Biologicals), mouse anti-tubulin (1:20,000; Abcam), rabbit anti-α2δ2 (1:5000; Alomone Labs, Jerusalem, Israel), mouse anti-ribosomal S6 (1:1000, Cell Signaling), rabbit anti-phospho-S6 (1:1000, Cell Signaling), and mouse anti-Ca_V_1.2 (1:2000; Neuromab). Antibody-bound membranes were incubated in fluorescence-conjugated secondary antibodies (AF680, Life Technologies; AF800, LiCor; 1:5000). Fluorescent images of the membranes were obtained using the Odyssey CLx infrared imaging system. Densitometry analyses of proteins were conducted using ImageJ (National Institutes of Health) software.

## Acknowledgments

This study was supported by National Institutes of Health NINDS NS105005 (KRG); the National Science Foundation IOS 1026527 and IOS 1355158 (KRG), Postdoctoral Research Fellowship in Biology DBI-1306528 (FN) and DBI-1103738 (LPC); Alzheimer’s Association AARF-19-614794 (FN); Department of Defense, United States Army Medical Research and Materiel Command USAMRMC Award W81XWH-14-1-0061 and W81XWH-19-1-0202 (KRG). National Institute of Aging Wake Forest School of Medicine Alzheimer’s Disease Research Center Pilot Grant P30AG049638 (KRG)

## Author contributions

FN, LPC, AU, SC, CDK, TM and KRG designed research. FN, LPC, AU, SN, CR, JP and WCT conducted experiments and analyzed data. CDK and SC provided the human samples. FN, LPC, AU, SN, and KRG wrote the manuscript.

